# Plasticity promotes persistence across novel environments in experimental microcosms

**DOI:** 10.1101/2025.11.01.686023

**Authors:** Emily A. Harmon, Charlex Malum, Sidharth Siddapureddy, David W. Pfennig

## Abstract

Phenotypic plasticity––the ability of organisms to adjust their traits in response to changes in their environment––has long been thought to prevent extinction in novel or changing environments. However, there are few tests linking plasticity to population persistence. Here, we show that plasticity promotes population persistence in replicate populations of rotifers. We experimentally exposed 33 clonal lines that varied in plasticity to over 20 novel environments for up to 40 overlapping generations. We found that clonal populations expressing plasticity in morphology and life history traits persisted longer and were less likely to go extinct in novel environments than populations not expressing such plasticity. These results directly link plasticity to population persistence in novel environments and suggest that plasticity can buy time for organisms in a changing world.

**Teaser:** A multigenerational experiment finds that environmentally induced traits allow populations to persist in novel environments.

## Introduction

When environments change faster than populations can adapt, those populations often go extinct (*1*). However, many organisms can respond immediately to such rapid environmental changes via phenotypic plasticity––the ability of a genotype or an individual organism to modify its phenotype in response to changes in its environment. Individuals expressing adaptive phenotypic plasticity in a new environment match their traits to current environmental conditions, survive, and proliferate, thereby reducing the risk that their population will go extinct (*2–8*). Given current profound and ongoing global changes precipitating a biodiversity crisis (*9*), a better understanding of this buffering capacity of plasticity is critical to predict which lineages and populations risk extinction.

The idea that plasticity promotes persistence in new environments is well supported by theory (*8, 10–15*) and consistent with evidence from natural populations and experimental work (*16–36*). For instance, plasticity in reproductive effort appears to have allowed dark-eyed juncos to persist in a new habitat (*16*), and, across bird species, those with innovative behavioral plasticity are less at risk of extinction (*18*). However, in many cases, it is difficult to show that plasticity directly contributes to population persistence. This requires not just demonstrating adaptive plasticity in a new environment but linking plasticity to population persistence across generations (*6, 37*). Further, few studies have demonstrated that plasticity was present before the invasion or environmental change occurred rather than evolving afterward.

Therefore, although the idea that plasticity can lessen extinction risk has existed for over a century under various guises––as components of the Baldwin Effect, the plasticity-mediated persistence hypothesis, or recently, the ‘buying time’ hypothesis (*6, 38, 39*)––a direct link between plasticity and persistence in new environments has not previously been established. This requires associating variation in plasticity expressed in a novel environment with variation in population persistence. The most powerful test would evaluate if experimental replicate populations that differ in degree of phenotypic plasticity vary in persistence in novel environments: more plastic populations should be more likely to persist than less plastic populations (*2, 6*). This approach would thereby determine if preexisting plasticity can prevent extinction by being beneficial in novel environments. Here, we undertake such an approach to investigate if plasticity protects populations from extinction.

We used a study system, rotifers, in which well-described preexisting plasticity can be induced across various laboratory environments and monitored for several generations. *Asplanchna brightwellii* are relatively large planktonic rotifers (up to 1 mm body length) found in freshwater worldwide and are important predators of cladocerans, copepods, ciliates, and other rotifers (*40, 41*). Clonal lines with a doubling time of about 20 hours can be easily maintained in the lab. Depending on intergenerational exposure to vitamin E gained from eating photosynthetic organisms or herbivorous prey, clones plastically vary in form (*i.e.,* morph) (*42–44*). This discrete plasticity results in individuals developing into either a default ‘alpha morph’ or a ‘beta morph’ if their mother consumed vitamin E (Fig. 1a) (*43, 45*). The larger beta morph is suited to conditions where large, herbivorous prey are abundant, and is characterized by body-wall outgrowths, or humps, that likely protect against cannibalism by other beta individuals (*46, 47*).

**Fig. 1.**
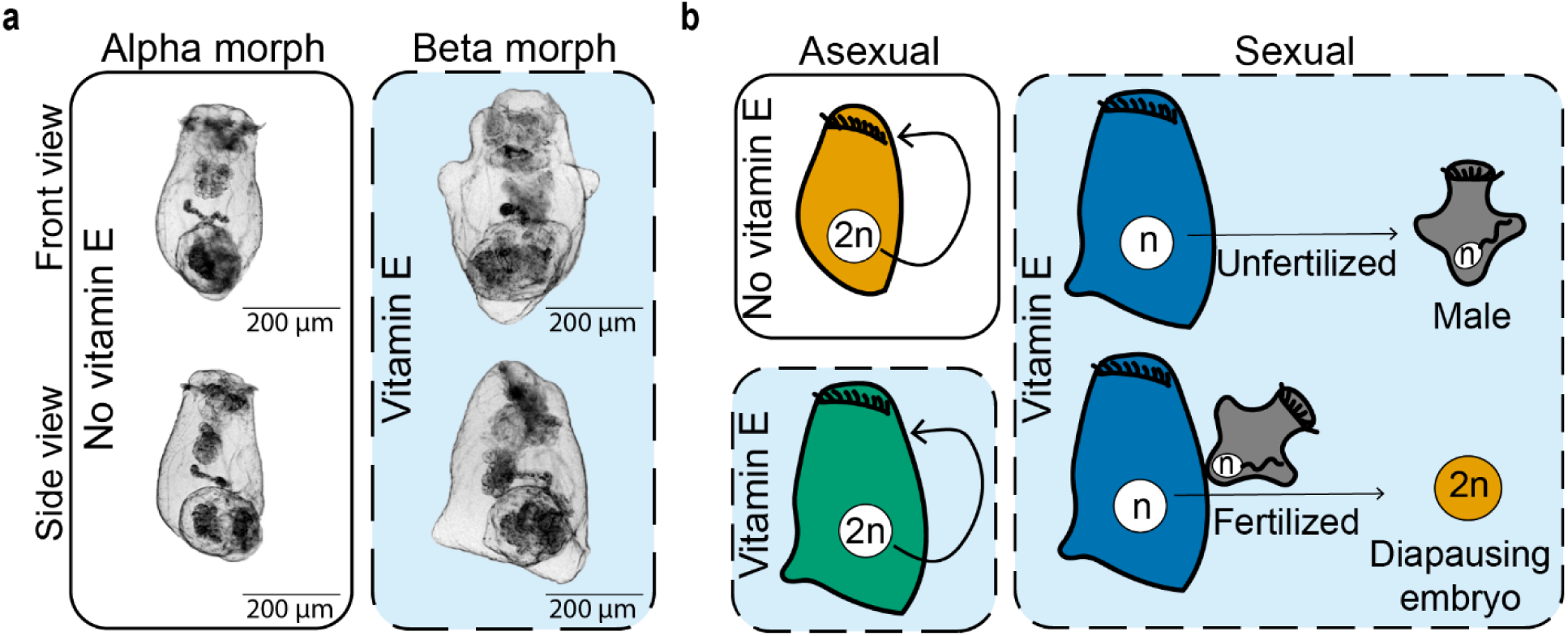
Morphological and reproductive plasticity in *A. brightwellii*. Vitamin E exposure *in utero* (dashed blue boxes) is a cue for the two forms of plasticity, which is fixed upon birth. (**a**) Stem females of a clonal line are the smaller (350-650 µm), sac-shaped alpha morph. Maternal ingestion of vitamin E induces successive generations of the larger (450-1000 µm), humped beta morph. (**b**) Alpha morphs (orange) and some beta morphs (green) reproduce asexually. Asexual females produce diploid eggs (white circles) that become females without fertilization. Some beta morphs reproduce sexually (blue); unfertilized haploid eggs develop into males (gray), and fertilized eggs develop into embryos within a cyst. These cysts have an extended diapause period before hatching as the stem female of a new clonal line.

In addition to plasticity in morphology, *A. brightwellii* exhibit plasticity in reproductive mode. Instead of the predominant mode of asexual reproduction (in which females have female offspring via diploid parthenogenesis), some beta morphs reproduce sexually, producing haploid male offspring or diploid diapausing embryos (Fig. 1b) (*48*). Diapausing embryos can withstand extreme environmental conditions such as pond drying, thereby allowing populations to avoid unsuitable environments. The proportion and degree of alpha, beta, and sexual morphotypes produced within a clonal line vary with maternal dietary vitamin E.

To ensure that *A. brightwellii* is appropriate to test for plasticity-mediated persistence, we first established that morphotype plasticity in *A. brightwellii* impacts fitness (and therefore could allow populations to maintain high fitness in new environments). Next, we exposed dozens of clonal lines of *A. brightwellii* to novel environments to evaluate if there was variable expression of preexisting plasticity in component traits of morphotype. Because the induced morph (the beta morph) can be either asexual or sexual, we could separate the role of plasticity and sexual reproduction in the response to the new environments. We measured responses to over 20 environments, treating each as a replicate new environment, which allowed for greater generalization of our findings.

We then tested for an association between variation in plasticity and variation in population persistence across the novel environments in two ways. First, we evaluated whether there was an overall relationship for clonal populations between persistence and the degree of plasticity (relative to control conditions) expressed in an environment. Second, we evaluated whether, more generally, plastic phenotypes – measured as trait variability across all the new and ancestral environments – were less likely to go extinct. If so, this would indicate that genotypes with greater plasticity across new and ancestral environments are better suited to escaping extinction in new and changing environments. Alternatively, plasticity-mediated persistence may be more context-dependent, with no genotype showing overall higher plasticity or extinction risk. This could arise through context-dependent plasticity: genotypes express alternate phenotypes only in a narrow range of environments, but their plasticity could be fortuitously beneficial in some novel environments. By evaluating multiple lineages across multiple environments, we can test not only for an association between plasticity and persistence but also characterize whether any plasticity-mediated persistence is a general property of highly plastic genotypes or context-dependent.

## Results

### Preexisting plasticity impacted fitness

We assessed whether morphotype plasticity in *A. brightwellii* is relevant for fitness under control laboratory conditions. We tracked the reproductive schedule from birth to death of 72 individuals representing 12 clonal lines to compare lifespan and fecundity of alpha morphs, asexual beta morphs, and sexual beta morphs (Supplemental Figure 1).

In typical laboratory conditions, alpha morphs lived an average of 1.5 times longer than the environmentally induced beta morphs (Fig. 2; linear mixed model [LMM] lifespan F_2,66_ = 8.07, p = 0.00073; Supplemental Tables 1-2). Alpha morphs also had more offspring than beta morphs (negative binomial generalized linear mixed model [GLMM] LRT = 9.58, df = 2, p= 0.0083; Supplemental Tables 1-2). On average, alpha morphs had 13.9 female offspring, asexual beta morphs had 9.2 female offspring, and sexual beta morphs had 4.6 female offspring (diapausing embryos), or if unfertilized, 11.3 male offspring (Table 1). Given this variation in fitness and life history strategy, the morph expressed by an individual in a novel environment could greatly impact its fitness and influence the persistence of its lineage.

**Fig. 2.**
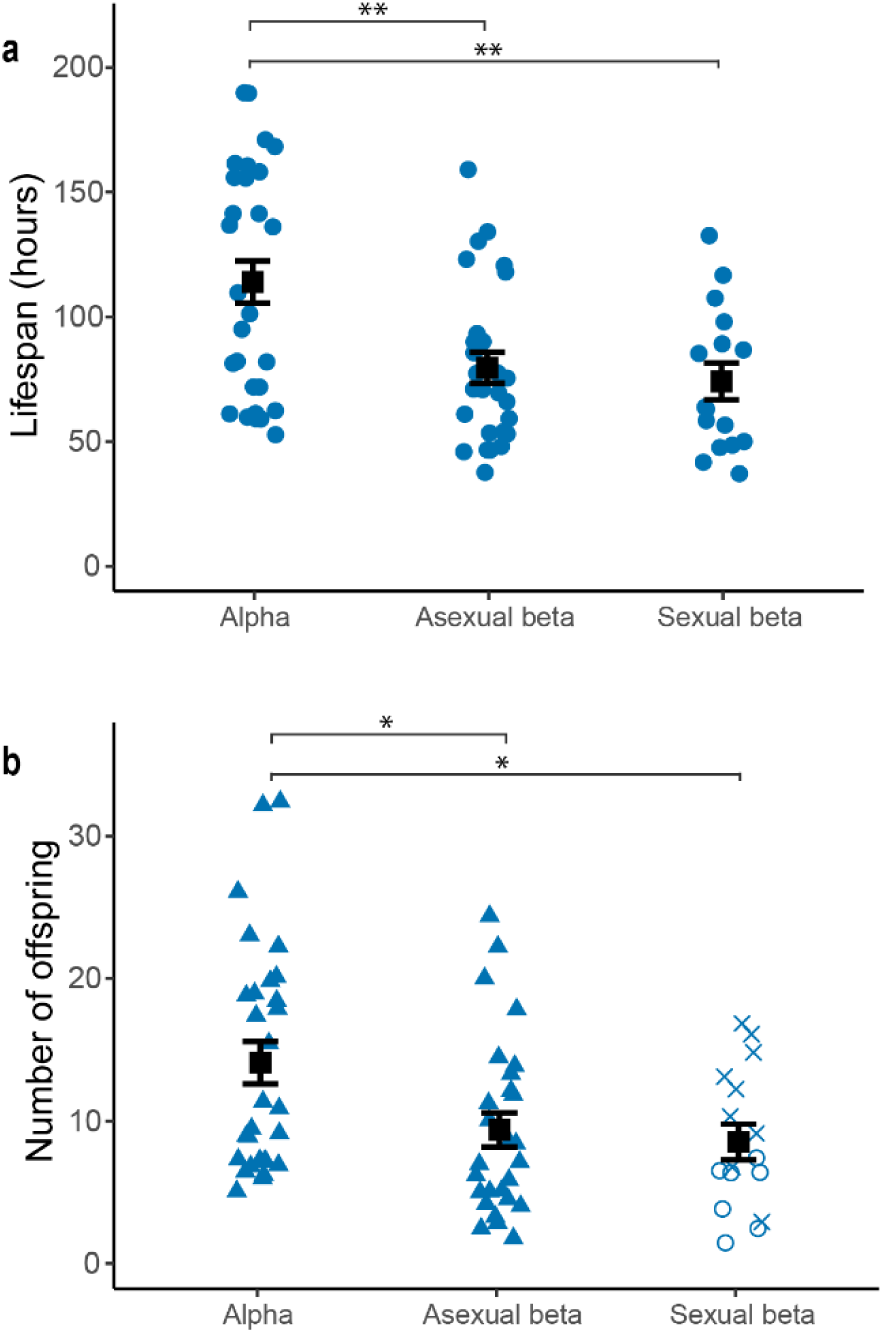
Preexisting plasticity impacts fitness. **(a)** Lifespan and **(b)** fecundity of alpha morphs (N = 29), asexual beta morphs (N = 27), and sexual beta morphs (N = 16). Values for individual rotifers in blue, morph mean ± 1 SEM in black. Offspring type shown by shape: female (blue closed triangle), diapausing embryo (blue open circle), male (blue cross). *p < 0.05, **p < 0.01 by Tukey post-hoc test (Supplemental Table 2).

**Table 1.** Lifetable and demographic parameters for morphs of *A. brightwellii* in the control environment.

| Morph | $R_0$ | G (hours) | r | Mean age at first birth (hours) |
| --- | --- | --- | --- | --- |
| Alpha | 13.9 | 75.4 | 0.035 | 33.9 |
| Asexual beta | 9.22 | 58.7 | 0.038 | 28.9 |
| Sexual beta | 4.57 | - | - | 41.1 |

Note: Not all values were calculated for sexual beta morphs because sexually produced offspring have an environmentally variable obligate period of diapause. $R_0$ , net reproductive rate: average number of female offspring a female will produce in her lifetime. G, mean generation time. r, intrinsic rate of increase.

**Table 2.** Environmental treatments. The baseline novel environments were elaborated upon to create the other novel environment treatments. The first set of experiments was housed in 24-well plates and the second set of experiments was housed in 50 mL vials as noted below.

| Environment Type | Container | Description |
| --- | --- | --- |
| Baseline | Well plate | 1 mL cultures were kept at room temperature and fed 500 paramecia/mL every other day |
|  | Vial | 20 mL cultures were kept at 20°C and fed 100 paramecia/mL every other day |
| Prey | Vial | Cultures were fed 5 *Lecane monostyla* */mL instead of paramecia |
|  | Vial | Cultures were fed 1 *Philodina* */mL instead of paramecia |
|  | Well plate | Cultures were fed 10 *Lepadella* */mL instead of paramecia |
| Competition | Vial | 5 *Didinium*<sup>†</sup> were added to cultures and replenished as needed |
|  | Well plate | Cultures were started with 10 rather than 5 *A. brightwellii* |
|  | Vial | Cultures were started with 10 rather than 5 *A. brightwellii* |
|  | Vial | 1 planarian was added to each culture and replenished as needed |
| Predator | Well plate | A Pasteur pipette was used to remove rotifers smaller than 400 µm every other day from cultures |
|  | Vial | Half of a culture was pipetted out through a 300 µm mesh, discarded, and replaced with EPA media every other day |
|  | Well plate | A Pasteur pipette was used to remove rotifers larger than 800 µm every other day from cultures |
|  | Vial | Half of a culture was pipetted out through a 300 µm mesh, the remnant was discarded, the filtered portion replaced and topped off with EPA media every other day |
|  | Well plate | Two copepods were added to cultures and replenished as needed |
| Temperature | Vial | Cultures were kept in an incubator at 15°C |
|  | Well plate | Cultures were kept in an incubator at 16°C |
|  | Well plate | Cultures were kept in an incubator at 30°C |
|  | Vial | Cultures were kept in an incubator at 25°C |
| Movement | Well plate | Cultures in well plates were kept on a rocker |
|  | Vial | Culture tubes were closed and kept laid down on a rocker |
| Water quality | Vial | Citric acid was used to maintain a pH of 6 |
|  | Vial | Miracle Gro® was used to maintain 10 mg/L total nitrogen |
|  | Well plate | Instant Ocean® was used to maintain 2 ppt salinity |
|  | Vial | Instant Ocean® was used to maintain 2 ppt salinity |
|  | Well plate | Resuspended fine sediment from a dried pond basin in AZ, USA was autoclaved and added to each culture and resuspended as needed |

\*other species of rotifers; <sup>†</sup>unicellular ciliates that, like *A. brightwellii*, feed on paramecia

### Replicate clonal populations differed in the degree of plasticity expressed in new environments

We evaluated whether the preexisting plastic ability to produce different morphotypes was variably expressed across new environments. We ran two sets of experiments varying in clonal lines and new environments used. The first exposed 19 clonal lines to 10 new environments and a control, and the second exposed 14 clonal lines to 13 new environments and a control. All environments contained trace amounts of vitamin E (*45, 49*) and were artificial lab environments consisting of small containers, high population densities, and a less diverse biological community than in the wild (*50*). The 23 new environments were chosen to be ecologically relevant and differentially select on morphology via changes in predation risk, feeding, and movement (Table 2). Each population was founded by five alpha morph clonemates and replicated three times per new environment (or fewer replicates if stock population sizes were limited).

We sampled approximately five rotifers from each clonal population after six days to assess their initial plastic responses to the new versus control environments. We measured the body size, shape, and reproductive mode as the component traits of morphotype plasticity. Each population in a new environment was given a context-dependent plasticity score for each trait calculated as the absolute value of the difference in traits produced in that population versus the traits produced by their clonemates in a control environment (Supplemental Figures 2-4). Note that despite the presence of sexual rotifers, all rotifers in our study were the product of asexual reproduction; diapausing individuals were not allowed to hatch, so sexual reproduction *per se* did not impact the outcome of our study (see Methods).

We confirmed that clonal lines varied in the expression of morphotype plasticity in the new environments. Indeed, we found that context-dependent plasticity in size (Fig. 3a; LMM: F_32,552_ = 2.82, p < 0.0001), shape (Fig. 3b; LMM: F_32,553_ = 4.23, p < 0.0001), and reproductive mode (Fig. 3c; LMM: F_32,552_ = 2.62, p < 0.0001) varied across clonal lines. We therefore successfully created a scenario in which replicate populations differ in degree of plasticity across novel environments.

**Fig. 3.**
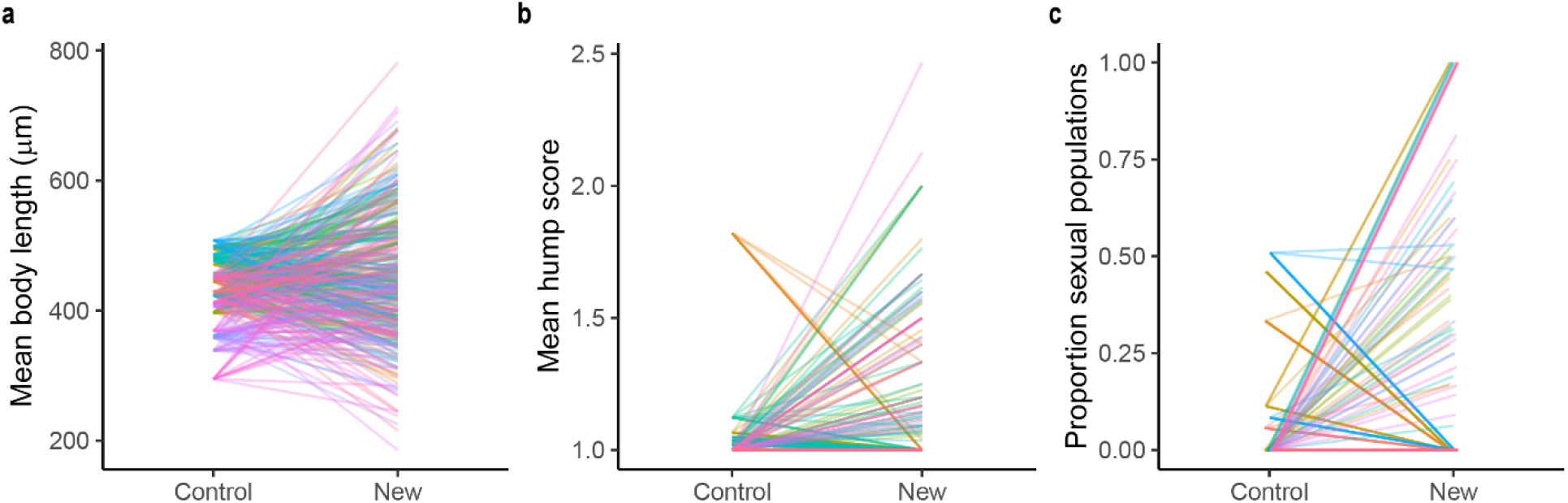
Replicate clonal populations differed in the degree of plasticity expressed in new environments. Clonal lines varied in plasticity of **(a)** body length (LMM; p < 0.0001), **(b)** body shape as measured by hump score (LMM; p < 0.0001), and **(c)** reproductive mode as measured by the proportion of the populations that reproduced sexually (LMM; p < 0.0001) expressed across new environments. Each line represents the traits produced by a clonal line in the control environment versus a new environment (see Supplemental Figures 2-4 for individual environments). Lines are colored by clonal line and transparent to better show overlapping patterns. For the first biological replicate, on average control N = 5 rotifers x 5 replicates, new environment N = 5 rotifers × 10 replicate new environments. For the second biological replicate, on average control N = 5 rotifers x 3 replicates, new environment N = 5 rotifers x 13 replicate new environments x 3 technical replicates.

### Plasticity was associated with population persistence in new environments

We recorded whether clonal populations persisted (N = 252) or not (N = 359) and how long they persisted for up to 50 days (19-40 overlapping generations; Table 1). To be conservative, we initially grouped populations that went dormant (no active culture but resting eggs present; N = 122) with the extinct populations (however, the main results were unchanged when we categorized populations that went dormant as persisting; see below). We asked whether variation in plasticity predicted variation in persistence. To do so, we fit GLMM and Cox mixed models that accounted for variance in extinction across clonal lines, spatial and temporal blocks, and environments (Supplemental Table 3).

We found that plasticity in reproductive mode predicted the *likelihood* of persistence (Supplemental Table 3). Specifically, the clonal populations most likely to persist were those expressing a greater degree of plasticity in shape (GLMM: LRT = 6.52, df = 1, p = 0.011; Figure 4a) and reproductive mode (GLMM: LRT = 4.39, df = 1, p = 0.036; Figure 4b). Similarly, plasticity predicted the *duration* of persistence (Supplemental Table 3). Increased plasticity in both shape and reproductive mode was associated with greater persistence. Each unit increase in standardized shape plasticity was associated with 42% less risk of extinction (hazard ratio [95% confidence interval] = 0.58 [0.41– 0.75], LRT = 10.73, p = 0.0011; Fig. 4c). Each unit increase in standardized reproductive mode plasticity was associated with 31% less risk of extinction (hazard ratio [95% confidence interval] = 0.69 [0.53– 0.85], LRT = 5.84, p = 0.016; Fig. 4d). In contrast to reproductive mode and shape, plasticity in body size was not an important predictor of population persistence (Supplemental Table 3).

**Fig. 4.**
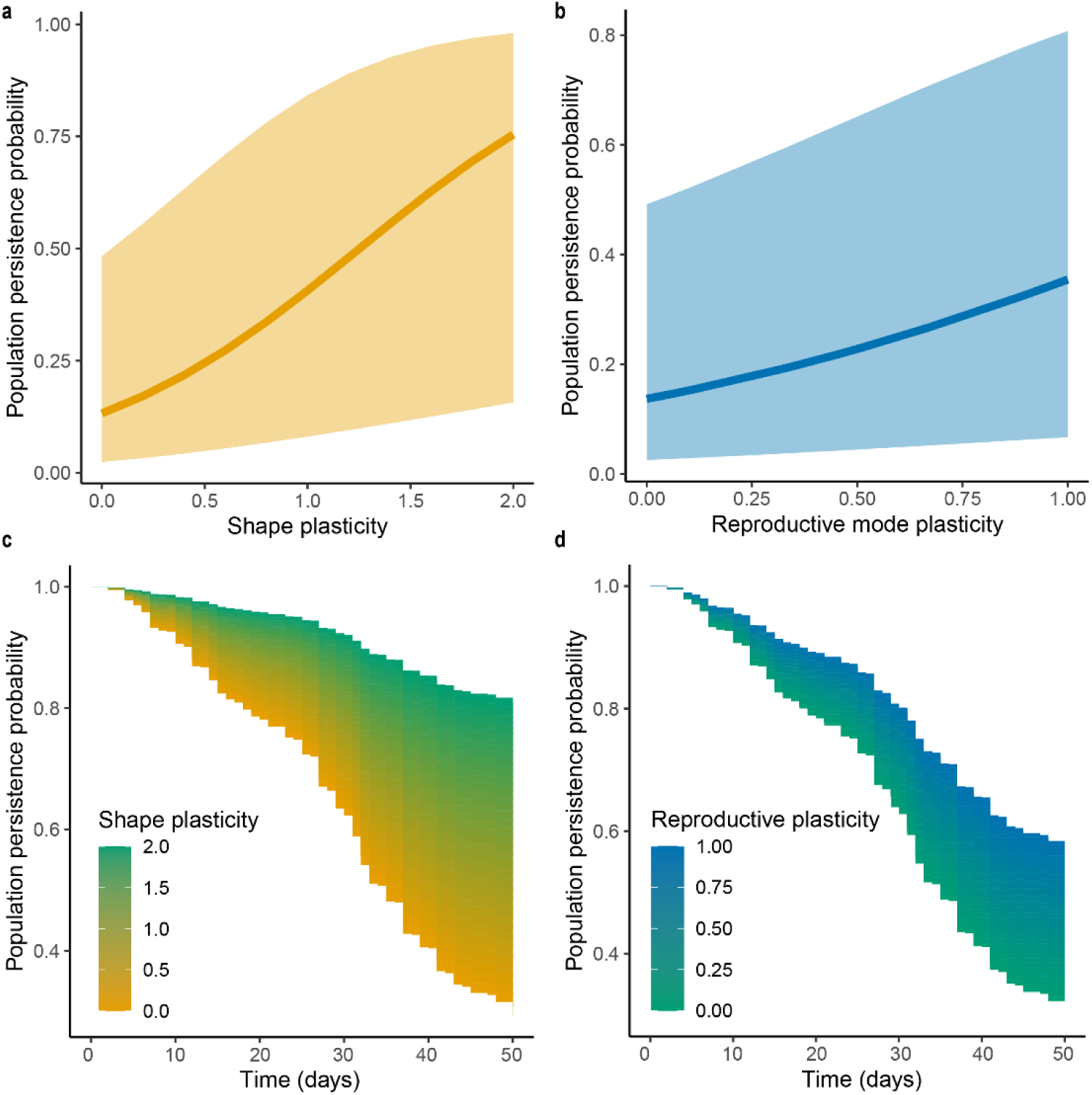
Plasticity is associated with population persistence in new environments. Probability of population persistence versus plasticity in shape (**a**) and reproductive mode (**b**). Lines show predicted values with 95% confidence interval of GLMM (Supplemental Table 3). Survival curve for populations colored by plasticity in shape (**c**) and reproductive mode (**d**). The curve is fit from a Cox mixed model (Supplemental Table 3). Plasticity is scored as the absolute value of the difference in traits produced by a clones in a new environment versus the control environment. We scored shape as the mean hump score and reproductive mode as the proportion of technical replicates that had any sexual reproduction (Fig. 3). N = 643 clonal populations.

These results are qualitatively similar if populations that went dormant are instead grouped with the persisting populations (Supplemental Table 4). Plasticity in shape and reproductive mode are associated with a lower likelihood of extinction and greater duration of persistence. In particular, reproductive mode plasticity (which allows for the creation of diapausing embryos) is associated with an even lesser risk of extinction if we assume that diapausing embryos could hatch in the future (hazard ratio [95% confidence interval] = 0.31 [0.04– 0.58], LRT = 18.66, p < 0.0001).

We found that variation in persistence in new environments was associated with variation in plasticity of shape and reproductive mode – the traits that determine morphotype. In sum, this finding is consistent with plasticity-mediated persistence: populations expressing greater plasticity in a new environment (relative to their original control environment) were more likely to persist in that environment.

#### General plasticity was not an indicator of persistence in new environments

Because populations expressing greater plasticity in a new environment were more likely to persist in that environment (see above), we next asked whether a genotype’s general plasticity across environments tended to predict its performance. We first measured general plasticity as the variance in each trait expressed across all the new environments. General plasticity varied across clonal lines in body size and shape (Levene’s test; size: F_32, 3843_ = 8.35, p < 0.0001; shape: F_32, 3842_ = 6.82, p < 0.0001) but not reproductive mode (F_32, 831_ = 0.92, p = 0.60).

We then evaluated whether clonal lines with a higher degree of general plasticity were more likely to persist in new environments. There was no relationship between general plasticity expressed across new environments and the likelihood a population went extinct (GLMM: size plasticity LRT = 0.41, df = 1, p = 0.52; shape plasticity LRT = 0.31, df = 1, p = 0.58; Supplemental Table 5) or duration of persistence (Cox mixed model: size plasticity LRT = 0.11, df = 1, p = 0.74; shape plasticity LRT = 0.22, df = 1, p = 0.41).

### Persistence of clonal lines in new environments was not predicted by ancestral plasticity

We next measured general plasticity as the degree of adaptive ancestral plasticity expressed by each genotype. We quantified plastic responses of the clonal lines to vitamin E by rearing clones in two environments for six days: without vitamin E and with enough vitamin E to elicit a robust morphological response in offspring (*42*). We measured the body length and hump score of individual rotifers and classified their reproductive mode (N = 1-7 clones from 3-5 replicate wells per clonal line per environment; Fig. 5). Each clonal line’s vitamin E plasticity was defined as the difference between environments in mean body size, mean hump score, and the percent of rotifers that were sexual (Supplemental Figure 5).

We found that the degree of beta expression under this strongly beta-inducing environment (*i.e.,* the preexisting plasticity to vitamin E), contrary to expectation, did not predict persistence in new environments. Clonal lines with more vitamin E plasticity were not more likely to persist (GLMM: vitamin E size plasticity LRT = 1.98, df = 1, p = 0.16; vitamin E shape plasticity LRT = 0.71, df = 1, p = 0.40; vitamin E reproductive mode plasticity LRT = 2.11, df = 1, p = 0.15; Supplemental Table 6) and did not persist longer across new environments (Cox mixed model: vitamin E size plasticity LRT = 0.047, df = 1, p = 0.83; vitamin E shape plasticity LRT = 0.050, df = 1, p = 0.82; vitamin E reproductive mode plasticity LRT = 0.045, df = 1, p = 0.83).

## Discussion

Testing the hypothesis that plasticity enables population persistence requires choosing replicate populations that differ in plasticity in a novel environment, and then assessing if variation in plasticity is associated with variation in persistence in new environments (*2, 6*). By using such an approach across 23 new environments, we found that morphotype plasticity in *A. brightwellii* was associated with lower extinction rates and longer persistence (Fig. 4). Specifically, populations expressing greater plasticity in a new environment were less likely to go extinct in that environment. Moreover, across different environments, different genotypes expressed high plasticity and persisted.

This study provides direct empirical support for plasticity-mediated persistence and sheds light on broad questions about the occurrence and consequences of plasticity-mediated persistence. Plasticity-mediated persistence can be interpreted in two main ways: either that generally plastic phenotypes should tend to persist in new environments, or that some lineages may express plasticity in a fortuitously adaptive manner in some new environments. Here, we find support for this latter, context-dependent form of plasticity-mediated persistence. In each environment, some lineages expressed preexisting plasticity to an extent or form that was adaptive – even if it could not do so across all the new environments. This aligns closely with the traditional evolutionary theory in which preexisting variation in a population may or may not be adaptive in a new setting. Plasticity, however, may promote persistence more than genetic variation because different phenotypes are expressed within a generation.

We may be primed to observe context-dependent plasticity-mediated persistence in systems such as *A. brightwellii* in which multiple morphotypes are produced (*i.e.,* a polyphenism). A polyphenism is typically thought of as a highly tailored response to specific environments in which alternate morphs would therefore be unlikely to be expressed, much less adaptive across many new environments (*15*). Other plastic traits, such as physiological and behavior responses, may be more primed to observe whether generally plastic phenotypes better avoid extinction across many environments. These traits tend to be more labile and responsive, and could serve a conserved, repurposed, or preadapted role across new environments (*51, 52*). However, despite the specificity of polyphenisms, we found that discrete plasticity in *A. brightwellii* can play adaptive roles across diverse new environments. It is possible, and even likely, that polyphenisms are underlain by multiple mechanisms of plasticity – including sensitive physiological and behavioral responses (*15, 37*).

Lineages expressing higher degrees of plasticity relative to the control environment tended to express more beta morphs in the new environments (Figure 3; most rotifers in control environments expressed the alpha morph). One major hypothesis for why the beta morph allowed for population persistence across new environments is that only beta morphs (not the alternative alpha morphs) can reproduce sexually. In theory, sexual reproduction should reduce extinction risk by increasing genetic variation and therefore the likelihood that adaptive variants can arise (*53–55*). However, we can reject this possibility. In our experiment, sexually produced offspring (diapausing embryos) were not allowed to contribute to active cultures. Consequently, populations producing sexual morphs did not benefit from increased genetic variation during the experiment. We also found no strong evidence across the new environments that populations varying in the expression of alpha and beta morphs had substantially different population dynamics (see Supplementary text). Any differences in fecundity among morphs may not be substantial in populations containing a mixture of morphs or may be counteracted by differences in survival in novel environments.

Therefore, the adaptive value of producing a beta morph across these new environments is likely due to an unmeasured trait that differs between the morphs but is generally beneficial across new environments. Indeed, many systems evolve polyphenisms through the genetic stabilization of phenotypes plastically produced in new (stressful) environments (*56–58*). The beta morph may then represent a stress-resistant morphotype with various physiological, behavioral, morphological, and life history adaptations that facilitate persistence in novel and changing environments.

Because plasticity mediated population persistence, these populations would presumably have greater temporal opportunity to evolve, enabling plasticity to facilitate evolution, albeit indirectly (*2*). The evolution of plasticity may be a common outcome of this ‘buying time’ hypothesis: genetic variation in plasticity can lift demographic constraints on evolution (*59*), leading to the evolution of higher adaptive plasticity within a population or evolutionary divergence between populations as less plastic genotypes are selected against. However, the buying time hypothesis is agnostic about the target of selection: such evolution might act on the plastic traits themselves, their degree of plasticity, or other traits (*2, 6*). Indeed, we observed changes within clonal lines during our experiment – some clonal populations that were purely asexual after initial exposure to new environments produced sexual morphs in later generations, suggesting that these populations evolved over the course of our experiments.

To better understand the evolutionary potential of plasticity-mediated persistence, future studies should monitor population sizes over time to evaluate how many generations plasticity ‘buys time.’ Future work can also evaluate how the dynamics of plasticity-mediated persistence change in populations that are not monoclonal, in which strains varying in plasticity must compete over multiple generations (*6*). Further investigations into environments with varying temporal dynamics (i.e., intensifying environments, fluctuating, predictable, or unpredictable) will also elucidate the bounds on when preexisting plasticity can be adaptively expressed in a new environment. Given the increasing appreciation for plasticity’s role in evolution, further work is required to determine whether and how plasticity-mediated persistence promotes evolution.

Rapid ongoing environmental change increasingly threatens many organisms and ecosystems globally, causing biodiversity loss and changing species distributions (*60*). Organisms must therefore respond to novel environments now more than ever, and evidence in nature suggests that plasticity facilitates coping with rapid environmental changes (*2, 61, 62*). Our results provide direct evidence that plasticity enables lineages to avoid extinction in novel or altered environments and suggest that plasticity *per se* could enable organisms to survive and persist in our rapidly changing world.

## Materials and Methods

### Collection

We established clonal lines of *A. brightwellii* by isolating stem females hatched from sexually produced diapausing embryos in pond sediment. The sediment came from ephemeral ponds in Texas, USA, provided by Dr. Elizabeth Walsh. From Hueco Tanks State Park & Historic Site, El Paso, TX, USA, we started seven clonal lines from Laguna Prieta (GPS coordinates: 31.9247, −106.0471) and ten clonal lines from Behind Ranch House playa (GPS coordinates: 31.9241, −106.0417). From the University of Texas at El Paso’s Indio Mountain Research Station, Hudspeth County, TX, USA, we started three clonal lines from Lonely Tank (GPS coordinates: 30.72787, −104.972), six clonal lines from Peccary Tank (GPS coordinates: 30.7609, −105.0043), and seven clonal lines from Rattlesnake Tank (GPS coordinates: 30.7505, −105.0075).

Other organisms were used as prey, competitors, and predators. We obtained *Paramecium aurelia*, *Didinium* (a unicellular ciliate that feeds primarily on paramecia), brown planarians, and rotifers *Philodina* sp. and *Lecane monostyla* from Carolina Biological Supply. We obtained *Lepadella* sp. rotifers from sediments collected from Carousel Park, Wilmington, DE, USA (GPS coordinates: 39.7307, −75.6856). Copepods were collected from Falls Lake, NC, USA (GPS coordinates: 36.0184, −78.7316).

### Culturing methods

We mass cultured *Paramecium aurelia* in 2 L Erlenmeyer flasks containing 1 L reconstituted moderate hard water (EPA media; 96 mg NaHCO_3_, 60 mg CaSO_4_,• 2H_2_O, 60 mg MgSO_4_, 4 mg KCl in 1 L distilled water (*63*)), 10 boiled wheat seeds, and approximately 1 g of nutritional yeast. Wheat seeds in prey media additionally acted as a source of trace amounts of vitamin E. We harvested cultures when they were about eight days old and filtered out biofilm through a 100 µm mesh. When necessary to obtain a concentrated solution, we concentrated paramecia on P8 grade filter paper and resuspended them in EPA media. Solution density was calculated from five sampled aliquots. A new culture was created and inoculated for each flask with 50-100 mL of an existing culture.

To start clonal lines of *A. brightwellii,* we rehydrated thin layers of pond sediment in 100 x 15 mm Petri dishes with EPA media at room temperature under UV lights. We checked the samples every 12 hours to isolate newly hatched stem females. We placed individual rotifers in 60 x 15 mm Petri dishes with a concentrated solution of paramecia until populations reached approximately ten individuals. At that point, they were split into two 120 mL jars containing 60 mL of paramecia solution. The clonal lines were cultured on paramecia at room temperature and ambient light for six months before use. Stock cultures of each clonal line were refreshed every four days by transferring 40 mL of the culture into a clean jar and bringing the volume to 60 mL with fresh EPA media. The remaining culture was discarded. Cultures were fed every other day with 15 mL of paramecia solution.

*Philodina*, *Lepadella*, and *L. monostyla* were separately cultured in 120 mL jars containing three boiled wheat seeds and EPA media. To harvest the rotifers, we centrifuged the culture medium at 500 x g for 10 minutes in conical tubes with a swinging bucket rotor. Concentrated rotifers were pipetted from the bottom of the tubes, and their density was calculated from five sampled aliquots. Stock cultures of *Didinium* were cultured in 120 mL jars of EPA media and fed approximately 15 mL *P. aurelia* every other day. Cultures were split into multiple jars as needed. Planarians were maintained in shallow glass bowls of EPA media covered in aluminum foil, fed a pea-sized amount of boiled egg yolk weekly, and given partial water changes as needed. Copepods were kept in 120 mL jars of EPA media and fed fish food *ad libitum*.

### Lifespan and fecundity of morphotypes

We evaluated differences in survival and reproduction for alpha morphs, asexual beta morphs, and sexual beta morphs across 12 clonal lines hatched from four ponds. Clonal lines were split into two 120 mL jars and exposed to either a control environment of EPA media or a vitamin E treatment (10^-7^ M *d*-α-tocopherol (*64*)) for five days to produce alpha and beta rotifers, respectively. We added 0.3 g/L penicillin to all solutions to curb excessive bacterial growth with vitamin E (*48*). Jars were fed equal amounts of paramecia every other day.

To obtain focal rotifers of varying morphotypes, we isolated three pairs of females from each jar and randomly distributed the pairs across wells of 24-well plates containing 1 mL of the source solution (control or vitamin E). We checked the wells every two hours until a female F1 neonate was produced (sexual F1 females could have been inseminated by F1 males during this time). We recorded the time of birth of the F1 female, isolated her in the well, and then fed the well approximately 500 paramecia. The F1 females were the focal rotifers for this experiment (N = 3 replicate clones x 3 clonal lines x 4 ponds x 2 environments = 72 F1 rotifers). Morphotype was recorded at maturity by visual inspection of shape and by offspring type (asexual morphs have female offspring, sexual morphs have male or diapausing offspring). Some focal rotifers from the control treatment developed as beta morphs, likely from low levels of vitamin E in prey media (*49*).

We kept the well plates at 20℃ and ambient light. Every other day, we used a Pasteur pipette to remove old media and waste which were replaced with new media and paramecia. Every 12 hours, we counted and removed F2 offspring born since the previous observation and recorded if the F1 rotifer had died. We continued until all F1 rotifers had died. We constructed a lifetable for each morph showing for each 12-hour age class how many F1 rotifers remained alive and how many F2 offspring were produced (Supplemental Figure 1). We used the standard formulae to calculate life history variables of survivorship, net reproductive rate, generation time, and intrinsic rate of increase (*65*).

### Plasticity and persistence in new environments

In two sets of experiments, we exposed clonal lines to new environments. In both experiments, we subsampled rotifers from each clonal population after six days (enough time for successive generations to mature; Table 1) to measure the expressed phenotypes. We tracked clonal populations in the new environments for approximately 50 days, designating them extinct when the active culture died out. This strict measure of extinction did not account for diapausing embryos, which were not allowed to hatch in the experiments. By excluding sexually produced diapausing embryos, there is limited potential for clonal lines to evolve in response to the new environments. We therefore can be more confident in the role of plasticity on persistence.

The first experimental set comprised 19 clonal lines established from four ponds. These clonal lines were reared in 10 new environments (Table 2 environments with container as “well plate”; N = 266 clonal populations in new environments) and a control environment (N = 5 replicates of each clonal line = 95 clonal populations in control environment). Baseline conditions included starting a population with five clonemates in 1 mL of EPA media in 24-well plates at room temperature. Clonal lines were randomly assigned to wells within a treatment block and well plates were kept in a stack at room temperature in ambient light. We removed waste and excess media from each well every other day using a Pasteur pipette under a dissecting microscope (to avoid removing *A. brightwellii*) and fed each well approximately 500 paramecia. We changed various aspects of these conditions for the other new environments (Table 2).

Each population was run in duplicate; the duplicate population was preserved in ethanol on day 6 to evaluate plastic responses across the new environments. Every five days we counted the rotifers of the remaining populations, removed diapausing embryos so that the culture remained clonal, and manually removed half the rotifers of each well (half the sum of females and diapausing embryos rounded down). By halving each population, we limited their ability to reach carrying capacity while preserving relative differences in population size among populations. We checked for extinction every other day and ran the experiment for 50 days.

The second experimental set comprised 14 newly established clonal lines from five ponds to limit adaptation to the lab environment. For this set, we modified the experimental procedures to be less labor intensive so that more replicates could be run simultaneously. We tested whether diapausing eggs hatch when left in culture, finding that 0 of 100 eggs hatched without first being desiccated and rehydrated. We therefore could leave the diapausing embryos in an experiment unit and still maintain a clonal culture. We scaled up the size of the containers to both achieve a higher carrying capacity (we do not attempt to control population size) and allow for more efficient methods of refreshing the water in a population. Finally, we decreased the baseline food from 500 to 100 paramecia/mL to increase competition.

The second set of clonal lines was exposed to 13 new environments and a control (Table 2 environments with container “vial”) with up to three replicates pending stock population size (N = 468 clonal populations in new environments; N = 36 clonal populations in control environment). Baseline conditions included allocating five clonemates to 20 mL of EPA media in a 50 mL centrifuge tube with a loose-fitting cap. Replicate tubes were arranged randomly in racks within an incubator at 20℃ and ambient light. Every other day, we fed the tubes 100 paramecia/mL, and every four days, we performed a partial water change through a 40 µm mesh to remove old media but not rotifers, added new EPA media, and restored the volume to 20 mL. Various aspects of this care varied across environments (Table 2).

Every other day, we checked whether each population persisted or had gone extinct. To assess plastic traits expressed in these new environments, we removed 1 mL of each population (5% of total volume) on the sixth day to preserve in ethanol for later phenotyping. We also recorded if there was evidence of sexual morphs in a population. We ran the experiment for 47 days.

The rotifers subsampled on day 6 from both sets of experiments were photographed using the 4x objective of a Leica compound microscope with the Olympus LC35 camera and 1x adapter using image acquisition program LCmicro V2.4. We measured the body length of each mature (gravid) rotifer in ImageJ 1.54g (*66*), scored body shape on a 1-3 scale (‘hump score’)(*45, 64*) and designated populations as sexual if any males or diapausing embryos were visible *in utero*.

### Ancestral plasticity and persistence

We tested the vitamin E-induced plasticity of each clonal line: 33 clonal lines sourced from five ponds in either a control environment of EPA media or in a vitamin E treatment of 10^-7^ M *d*-α-tocopherol (*64*). Three to five replicate clonal populations (depending on stock population density) of five clones were randomly distributed across the wells of 24-well plates, each containing 1 mL of control or vitamin E solution, 0.3 g/L penicillin to curb bacterial growth that occurs with exogenous vitamin E (*48*), and 500 paramecia/mL. Paramecia media may introduce further trace amounts of vitamin E to all wells. Well plates were kept at 20℃ and ambient light. We performed a partial water change every other day with a Pasteur pipette and fed each well 500 paramecia. After six days, we added 95% ethanol to each well to fix the rotifers.

We quantified body size, shape and reproductive mode for approximately five mature females from each well (on average N = 20 rotifers per clonal line per environment). Clonal lines with fewer than three mature female rotifers measured per environment due to extinction or population demographics were removed from analysis (N = 3 clonal lines). We photographed rotifers with an Olympus LC30 camera with 0.5x adapter on an Olympus CX43 microscope at 4x. We measured body length from these photos in ImageJ 1.54g (*66*). We classified the shape of each rotifer as humped or not. When possible (for 857 of 1214 rotifers), we determined reproductive mode from the sex of offspring *in utero*. We calculated the plasticity of each trait (body length, hump score, percent of population that is sexual) for each clonal line as the difference in values between the two environments.

### Statistical analysis

We conducted all analyses in R (*67*) (all raw data are available (*68*)). We tested the fixed effects of models by comparing the model of interest to a null model. We tested LMMs with a Kenward-Roger F test (function *drop1*), GLMMs with a likelihood ratio test (function *drop1*), and Cox mixed models with a type II analysis of variance (function *Anova* in the package ‘car’ (*69*)). Random effects accounting for no variance were dropped from models. Model assumptions throughout were verified from model diagnostic tests and plots from package ‘DHARMa’ (*70*). If assumptions of residual normality or homoskedasticity were violated, we bootstrapped confidence intervals for comparison (function *bootstrap* from the package ‘lmeresampler’ (*71*)).

#### Lifespan and fecundity of morphotypes

We evaluated whether lifespan varied across alpha morphs (N = 29), asexual beta morphs (N = 27), and sexual beta morphs (N = 16) via a LMM fitted with restricted maximum likelihood using morph as the fixed effect and clonal line as a random intercept (function *lmer* from the package ‘lme4’ (*72*); Supplemental Table 1). To model number of offspring per morph, we used a negative binomial GLMM with the package *glmmTMB* with morph as the fixed effect and clonal line as a random intercept. Pairwise comparisons of the estimated marginal means for morphs were performed using posthoc tests with the Tukey method for multiple comparison p-value adjustments (‘emmeans’ package (*73*)).

#### Variation in plasticity in new environments

We evaluated whether clonal lines varied in plasticity across the new environments. Populations that went extinct within the first six days or had samples with only immature rotifers or poor preservation quality were excluded from analysis (N = 98). For each clonal population, we calculated the average body length, average hump score, and categorized the presence or absence of sexual morphs (representing on average N = 5 clones measured per population). Each clonal population was given a plasticity score for body size, shape, and reproductive mode that is the absolute value of the difference between their traits and those expressed by contemporary clonemates in the control environment. To assess whether clones varied in plasticity across the environments, we fit LMMs with the plasticity measurement as the response variable, clonal line as a fixed effect, and environment, pond, and spatial block nested in temporal round as random intercepts.

#### Context-dependent plasticity and persistence

To evaluate whether variation in plasticity expressed across new environments was associated with persistence across environments, we generated models including fixed effects of the body size plasticity, body shape plasticity, reproductive mode plasticity, and temporal round. We standardized numerical predictor variables to a mean of 0 and standard variation of 0.5 to conduct model averaging and allow direct comparison of predictor effects (*74*). We did not include any interaction terms as we had no strong *a priori* expectations for any. We included random intercepts of clonal line nested in source pond, environment type, and spatial blocking (Supplemental Table 3). Any random effects accounting for no variance were dropped from the model.

We used GLMMs with a binomial distribution to evaluate the likelihood of persistence and mixed effects Cox models (package ‘coxme’ (*75*)) to evaluate the duration of persistence (N = 643 populations). For GLMMs, the response variable was whether the population persisted, and for the Cox mixed effect models, the response variable was the censored time to extinction. We evaluated variance inflation factors (VIF) to check for multicollinearity. We ran a set of models in which dormant populations were grouped with the extinct populations (Supplemental Table 3) and a set in which dormant populations were grouped with the persisting populations (Supplemental Table 4).

#### General plasticity and persistence

We measured the general plasticity in size, shape, and reproductive mode for a genotype as the variance across all tested environments. We used Levene’s tests to evaluate whether genotypes varied in general plasticity. We then analyzed whether general plasticity predicted a clonal line’s ability to avoid extinction across all the environments (N = 643 populations). We constructed GLMM and Cox mixed models using the general plasticity metrics as fixed effects and clonal line, treatment, and round nested in spatial block as random intercepts (Supplemental Table 5).

#### Ancestral plasticity and persistence

We analyzed whether vitamin E plasticity predicted a clonal line’s ability to persist in the new environments (N = 549 populations). As above, we constructed GLMM and Cox mixed models, here using vitamin E plasticity metrices as the fixed effects and clonal line, treatment, and round nested in spatial block as random intercepts (Supplemental Table 6).

## Acknowledgements

Sediment was collected from Hueco Tanks State Park and Historic Site (Texas) under permits 07-02 and 07-21. Sediment was collected from Indio Mountain Research Station by O. da Cunha. Many thanks to E.J. Walsh for providing sediment samples containing *A. brightwellii* diapausing embryos. We thank K. Chen, E. Gurkin, J. Kim, and T. Li for laboratory assistance. We are grateful to K. Pfennig, J. Kingsolver, A. Isdaner, P. Kelly, K. Hill, K. Sockman, B. Goldstein, W. Park, S. Oliveaux, and J. Drennan for discussion and comments.

## Author contributions

EAH and DWP conceptualized and designed the study. CM designed and ran the lifetable experiment. EAH and SS ran the persistence experiments. EAH analyzed the data and wrote the original draft. DWP and EAH reviewed and edited the paper. All authors reviewed and approved the final draft.

## Data and materials availability

The data and code generated in this work have been deposited at figshare and are publicly available as of the date of publication at https://doi.org/10.6084/m9.figshare.25870906.

## References

1. G. Bell, A. Gonzalez, Adaptation and evolutionary rescue in metapopulations experiencing environmental deterioration. Science 332, 1327–1330 (2011).

2. S. E. Diamond, R. A. Martin, “Buying time: plasticity and population persistence” in Phenotypic Plasticity and Evolution: Causes, Consequences, Controversies, D. W. Pfennig, Ed. (CRC Press, Boca Raton, FL, 2021), pp. 185–209.

3. M. J. West-Eberhard, Developmental Plasticity and Evolution (Oxford University Press, New York, NY, 2003).

4. C. D. Schlichting, M. A. Wund, Phenotypic plasticity and epigenetic marking: an assessment of evidence for genetic accommodation. Evolution 68, 656–672 (2014).

5. S. E. Diamond, R. A. Martin, The interplay between plasticity and evolution in response to human-induced environmental change. F1000Res 5, 2835 (2016).

6. M. R. Morris, Plasticity-mediated persistence in new and changing environments. Int. J. Evol. 2014, 416497 (2014).

7. R. J. Fox, J. M. Donelson, C. Schunter, T. Ravasi, J. D. Gaitan-Espitia, Beyond buying time: the role of plasticity in phenotypic adaptation to rapid environmental change. Philos. Trans. R. Soc. B: Biol. Sci. 374, 20180174 (2019).

8. S. M. Scheiner, M. Barfield, R. D. Holt, The genetics of phenotypic plasticity. XV. Genetic assimilation, the Baldwin effect, and evolutionary rescue. Ecol. Evol. 7, 8788–8803 (2017).

9. S. Díaz, J. Settele, E. S. Brondízio, H. T. Ngo, J. Agard, A. Arneth, P. Balvanera, K. A. Brauman, S. H. M. Butchart, K. M. A. Chan, L. A. Garibaldi, K. Ichii, J. Liu, S. M. Subramanian, G. F. Midgley, P. Miloslavich, Z. Molnár, D. Obura, A. Pfaff, S. Polasky, A. Purvis, J. Razzaque, B. Reyers, R. R. Chowdhury, Y.-J. Shin, I. Visseren-Hamakers, K. J. Willis, C. N. Zayas, Pervasive human-driven decline of life on Earth points to the need for transformative change. Science 366, eaax3100 (2019).

10. T. D. Price, A. Qvarnström, D. E. Irwin, The role of phenotypic plasticity in driving genetic evolution. Proc. R. Soc. B: Biol. Sci. 270, 1433–1440 (2003).

11. R. Lande, Adaptation to an extraordinary environment by evolution of phenotypic plasticity and genetic assimilation. J. Evol. Biol. 22, 1435–1446 (2009).

12. L. M. Chevin, R. Lande, G. M. Mace, Adaptation, plasticity, and extinction in a changing environment: towards a predictive theory. PLoS Biol. 8, e1000357 (2010).

13. J. Ashander, L.-M. Chevin, M. L. Baskett, Predicting evolutionary rescue via evolving plasticity in stochastic environments. Proc. R. Soc. B: Biol. Sci. 283, 20161690 (2016).

14. S. M. Scheiner, M. Barfield, R. D. Holt, The genetics of phenotypic plasticity. XVII. Response to climate change. Evol. Appl. 13, 388–399 (2020).

15. E. C. Snell-Rood, M. E. Kobiela, K. L. Sikkink, A. M. Shephard, Mechanisms of plastic rescue in novel environments. Annu. Rev. Ecol. Evol. Syst. 49, 331–354 (2018).

16. P. J. Yeh, T. D. Price, Adaptive phenotypic plasticity and the successful colonization of a novel environment. Am. Nat. 164, 531–542 (2004).

17. D. W. Pfennig, M. McGee, Resource polyphenism increases species richness: a test of the hypothesis. Philos. Trans. R. Soc. B: Biol. Sci. 365, 577–591 (2010).

18. S. Ducatez, D. Sol, F. Sayol, L. Lefebvre, Behavioural plasticity is associated with reduced extinction risk in birds. Nat. Ecol. Evol. 4, 788–793 (2020).

19. A. V. Badyaev, Evolutionary significance of phenotypic accommodation in novel environments: an empirical test of the Baldwin effect. Philos. Trans. R. Soc. B: Biol. Sci. 364, 1125–1141 (2009).

20. S. Volis, D. Ormanbekova, K. Yermekbayev, Role of phenotypic plasticity and population differentiation in adaptation to novel environmental conditions. Ecol. Evol. 5, 3818–3829 (2015).

21. J. Hua, D. K. Jones, B. M. Mattes, R. D. Cothran, R. A. Relyea, J. T. Hoverman, The contribution of phenotypic plasticity to the evolution of insecticide tolerance in amphibian populations. Evol. Appl. 8, 586–596 (2015).

22. A. Corl, K. Bi, C. Luke, A. S. Challa, A. J. Stern, B. Sinervo, R. Nielsen, The genetic basis of adaptation following plastic changes in coloration in a novel environment. Curr. Biol. 28, 2970–2977. e2977 (2018).

23. A. J. Garrison, L. A. Norwood, J. K. Conner, Plasticity-mediated persistence and subsequent local adaptation in a global agricultural weed. Evolution 78, 1804–1817 (2024).

24. T. D. Price, P. J. Yeh, B. Harr, Phenotypic plasticity and the evolution of a socially selected trait following colonization of a novel environment. Am. Nat. 172, S49–S62 (2008).

25. C. A. Handelsman, E. D. Broder, C. M. Dalton, E. W. Ruell, C. A. Myrick, D. N. Reznick, C. K. Ghalambor, Predator-induced phenotypic plasticity in metabolism and rate of growth: rapid adaptation to a novel environment. Integr. Comp. Biol. 53, 975–988 (2013).

26. D. Sol, Revisiting the cognitive buffer hypothesis for the evolution of large brains. Biol. Lett. 5, 130–133 (2009).

27. D. Sol, S. Timmermans, L. Lefebvre, Behavioural flexibility and invasion success in birds. Anim. Behav. 63, 495–502 (2002).

28. O. Vedder, S. Bouwhuis, B. C. Sheldon, Quantitative assessment of the importance of phenotypic plasticity in adaptation to climate change in wild bird populations. PLoS Biol. 11, e1001605 (2013).

29. A. P. Møller, D. Rubolini, E. Lehikoinen, Populations of migratory bird species that did not show a phenological response to climate change are declining. PNAS 105, 16195–16200 (2008).

30. A. M. Davidson, M. Jennions, A. B. Nicotra, Do invasive species show higher phenotypic plasticity than native species and, if so, is it adaptive? A meta-analysis. Ecol. Lett. 14, 419–431 (2011).

31. G. C. Trussell, L. D. Smith, Induced defenses in response to an invading crab predator: an explanation of historical and geographic phenotypic change. PNAS 97, 2123–2127 (2000).

32. S. Bonamour, L.-M. Chevin, A. Charmantier, C. Teplitsky, Phenotypic plasticity in response to climate change: the importance of cue variation. Philos. Trans. R. Soc. B: Biol. Sci. 374, 20180178 (2019).

33. A. Markov, S. Ivnitsky, Evolutionary role of phenotypic plasticity. Moscow University Biological Sciences Bulletin 71, 185–192 (2016).

34. J. M. Gómez, A. González-Megías, C. Armas, E. Narbona, L. Navarro, F. Perfectti, The role of phenotypic plasticity in shaping ecological networks. Ecol. Lett. 26, S47–S61 (2023).

35. C. T. Kremer, S. B. Fey, A. A. Arellano, D. A. Vasseur, Gradual plasticity alters population dynamics in variable environments: thermal acclimation in the green alga Chlamydomonas reinhartdii. Proc. R. Soc. B: Biol. Sci. 285, 20171942 (2018).

36. M. Rescan, D. Grulois, E. Ortega-Aboud, L.-M. Chevin, Phenotypic memory drives population growth and extinction risk in a noisy environment. *Nat*. Ecol. Evol. 4, 193–201 (2020).

37. A. P. Hendry, Key questions on the role of phenotypic plasticity in eco-evolutionary dynamics. J. Hered. 107, 25–41 (2016).

38. J. M. Baldwin, A new factor in evolution (Continued). Am. Nat. 30, 536–553 (1896).

39. E. Pennisi, Buying time. Science 362, 988–991 (2018).

40. E. J. Walsh, H. A. Smith, R. L. Wallace, Rotifers of temporary waters. Int. Rev. Hydrobiol. 99, 3–19 (2014).

41. C. Williamson, J. J. Gilbert, “Variation among zooplankton predators: the potential of *Asplanchna*, *Mesocyclops*, and *Cyclops* to attack, capture, and eat various rotifer prey” in Evolution and ecology of zooplankton communities, W. C. Kerfoot, Ed. (The University Press of New England, Hanover, London, 1980), pp. 509–517.

42. J. J. Gilbert, Non-genetic polymorphisms in rotifers: environmental and endogenous controls, development, and features for predictable or unpredictable environments. Biol. Rev. 92, 964–992 (2017).

43. C. W. Birky, Jr., Studies on the physiology and genetics of the rotifer, *Asplanchna*. I. Methods and physiology. J. Exp. Zool. 155, 273–291 (1964).

44. C. W. Birky, Jr., The developmental genetics of polymorphism in the rotifer *Asplanchna*. I. Dietary vitamin E control of mitosis and morphogenesis in embryos. J. Exp. Zool. 169, 205–210 (1968).

45. E. A. Harmon, D. W. Pfennig, Diet-induced polyphenism in *Asplanchna brightwellii* rotifers: developmental and evolutionary implications of trait variation and integration. Hydrobiologia, (2025).

46. J. J. Gilbert, Female polymorphism and sexual reproduction in the rotifer *Asplanchna*: evolution of their relationship and control by dietary tocopherol. Am. Nat. 116, 409–431 (1980).

47. J. J. Gilbert, The adaptive significance of polymorphism in the rotifer *Asplanchna*. Humps in males and females. Oecologia 13, 135–146 (1973).

48. J. J. Gilbert, Polymorphism and sexuality in the rotifer *Asplanchna*, with special reference to the effects of prey-type and clonal variation. Arch. Hydrobiol. 75, 442–483 (1975).

49. J. J. Gilbert, Dietary control of sexuality in the rotifer *Asplanchna brightwelli* Gosse. Physiol. Zool. 41, 14–43 (1968).

50. J. Conde-Porcuna, S. Declerck, Regulation of rotifer species by invertebrate predators in a hypertrophic lake: selective predation on egg-bearing females and induction of morphological defences. J. Plankton Res. 20, 605–618 (1998).

51. D. W. Pfennig, “Key questions about phenotypic plasticity” in Phenotypic plasticity & evolution: causes, consequences, controversies, D. W. Pfennig, Ed. (CRC Press, Boca Raton, FL, 2021), pp. 55–87.

52. S. E. Coates, A. A. Comeault, D. P. Wood, M. F. Fay, S. Creer, O. G. Osborne, L. T. Dunning, A. S. Papadopulos, Plastic responses to past environments shape adaptation to novel selection pressures. PNAS 122, e2409541122 (2025).

53. B. O. Bengtsson, Genetic variation in organisms with sexual and asexual reproduction. J. Evol. Biol. 16, 189–199 (2003).

54. J. Crow, An advantage of sexual reproduction in a rapidly changing environment. J. Hered. 83, 169–173 (1992).

55. J. Lachapelle, G. Bell, Evolutionary rescue of sexual and asexual populations in a deteriorating environment. Evolution 66, 3508–3518 (2012).

56. A. V. Badyaev, Stress-induced variation in evolution: from behavioural plasticity to genetic assimilation. Proc. R. Soc. B: Biol. Sci. 272, 877–886 (2005).

57. A. M. Shapiro, “Seasonal polyphenism” in Evolutionary Biology*: Volume* 9, M. K. Hecht, W. C. Steere, B. Wallace, Eds. (Springer, 1976), pp. 259–333.

58. Y. Suzuki, H. F. Nijhout, Evolution of a polyphenism by genetic accommodation. Science 311, 650–652 (2006).

59. R. Gomulkiewicz, D. Houle, Demographic and genetic constraints on evolution. Am. Nat. 174, E218–E229 (2009).

60. C. Parmesan, G. Yohe, A globally coherent fingerprint of climate change impacts across natural systems. Nature 421, 37–42 (2003).

61. M. J. West-Eberhard, Phenotypic plasticity and the origins of diversity. Annu. Rev. Ecol. Syst. 20, 249–278 (1989).

62. E. A. Harmon, D. W. Pfennig, Evolutionary rescue via transgenerational plasticity: evidence and implications for conservation. Evol. Dev. 23, 292–307 (2021).

63. W. H. Peltier, C. I. Weber, “Methods for measuring the acute toxicity of effluents to freshwater and marine organisms” (EPA/600 & 4-85/013, US Environmental Protection Agency, 1985).

64. C. W. Birky, Jr., The developmental genetics of polymorphism in the rotifer *Asplanchna*. III. Quantitative modification of developmental responses to vitamin E, by the genome, physiological state, and population density of responding females. J. Exp. Zool. 170, 437–448 (1969).

65. C. J. Krebs, Ecology: The Experimental Analysis of Distribution and Abundance (Harper and Row, New York, NY, 1985).

66. C. A. Schneider, W. S. Rasband, K. W. Eliceiri, NIH Image to ImageJ: 25 years of image analysis. Nat. Methods 9, 671–675 (2012).

67. R Core Team, R: a language and environment for statistical computing. https://www.R-project.org/ (2024).

68. E. A. Harmon, C. Malum, S. Siddapureddy, D. W. Pfennig, Data for: Plasticity promotes population persistence. figshare, (2024).

69. J. Fox, S. Weisberg, D. Adler, D. Bates, G. Baud-Bovy, S. Ellison, D. Firth, M. Friendly, G. Gorjanc, S. Graves, Package ‘car’. Vienna: R Foundation for Statistical Computing 16, (2012).

70. F. Hartig, DHARMa: Residual Diagnostics for Hierarchical (Multi-Level / Mixed) Regression Models. https://CRAN.R-project.org/package=DHARMa (2022).

71. A. Loy, S. Steele, J. Korobova, lmeresampler: Bootstrap Methods for Nested Linear Mixed-Effects Models. https://CRAN.R-project.org/package=lmeresampler (2023).

72. D. Bates, M. Mächler, B. Bolker, S. Walker, Fitting linear mixed-effects models using lme4. J. Stat. Softw. 67, 1–48 (2015).

73. R. V. Lenth, emmeans: estimated marginal means, aka least-squares means. https://CRAN.R-project.org/package=emmeans (2024).

74. C. E. Grueber, S. Nakagawa, R. J. Laws, I. G. Jamieson, Multimodel inference in ecology and evolution: challenges and solutions. J. Evol. Biol. 24, 699–711 (2011).

75. T. M. Therneau, coxme: Mixed Effects Cox Models. https://CRAN.R-project.org/package=coxme (2022).

