## Supplementary material for "Plasticity promotes persistence across novel environments in experimental microcosms": Document S1

Science Advances

AAAS

Emily Harmon *et al.*

**This PDF file includes:**

Supplementary Text
Figs. S1 to S5
Tables S1 to S9

**Supplementary Text**

It is possible that populations expressing beta morphs persist better in new environments because of different population dynamics associated with differences in morph life histories. Because sexual morphs suffer the two-fold cost of producing males and diapausing embryos, we might predict that these populations grew more slowly than populations of alpha morphs (which produce more offspring under control conditions; Fig. 2). Cannibalism of beta morphs on smaller rotifers may also slow population growth. If beta morphs also have fewer offspring in the new environments, this slower population growth might make populations more stable in small experimental microcosms, avoiding population crashes possible with an overshoot of carrying capacity. There is also some evidence in *Asplanchna* that the production of sexual morphs increases with population density.

In the first round of the experiment in which rotifers were exposed to new environments, we recorded and manually halved population size every five days. To evaluate if higher population density may drive formation of the sexual beta morph, we classified the overall morphotype of clonal populations sampled on day 6 based on their shape and reproductive mode (alpha morphs have hump score = 1 and are asexual; beta morphs have hump score > 1 and/or are sexual). Six days was chosen to be confident that the rotifers measured in the environment were subject to that new environment for their entire embryonic development (Table 1). We then used binomial GLMMs with (1) the proportion of beta morphs and (2) the proportion of beta morphs that were sexual as response variables, population size at day 5 as the fixed effect, and environment, clonal line nested in pond, and rack as random effects (random effects accounting for no variance were dropped from the models; Supplemental Table 6). We tested fixed effects with likelihood ratio tests. There was no evidence that population density on day five predicted the production of beta morphs (which were assayed the next day; GLMM LRT = 0.059, df = 1, $P$ = 0.81), and the proportion of populations with beta morphs that could reproduce sexually was also not associated with previous population density (GLMM LRT = 0.083, df = 1, $P$ = 0.77; Supplemental Table 7).

Further, there were no humped or sexual morphs induced in environments with double the population density (Supplemental Figures 3-4 “crowd” environments).

To evaluate if populations with beta morphs tended to grow more slowly than populations with alpha morphs, we tested for a relationship between the presence of beta morphs as measured on day 6 and the rate of population increase between days 5-10. We calculated the intrinsic rate of increase as $r = (\ln(N_t) - \ln(N_0))/t$. We used a LMM and Kenward-Roger F test to evaluate if the fixed effect of morph predicted r. Environment, pond, and rack were used as random effects (clonal line accounted for no variance; Supplemental Table 8). The rate of population growth between days 5-10 was unrelated to morphotype measured in the population on day 6 (LMM: $F_{1,195} = 1.39$, $P = 0.24$; Supplemental Table 8). Similarly, the stability of populations across the timeseries (absolute value of r at each timestep) was unrelated to the plastic morphotype (LMM: F1, 250 = 2.84, $P = 0.093$; Supplemental Table 9).

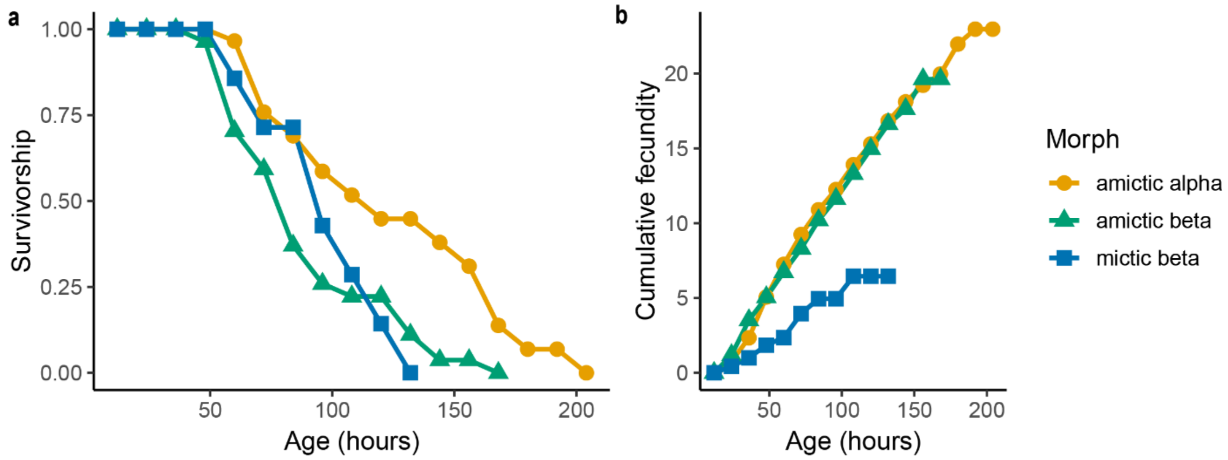


**Fig. S1. Lifetable results for different morphs.** Survivorship (**a**) and cumulative fecundity (**b**) for cohorts of alpha morphs (N = 29), asexual (amictic) beta morphs (N = 27), and sexual (mictic) beta morphs (N = 16). Comparative statistics in Figure 2.

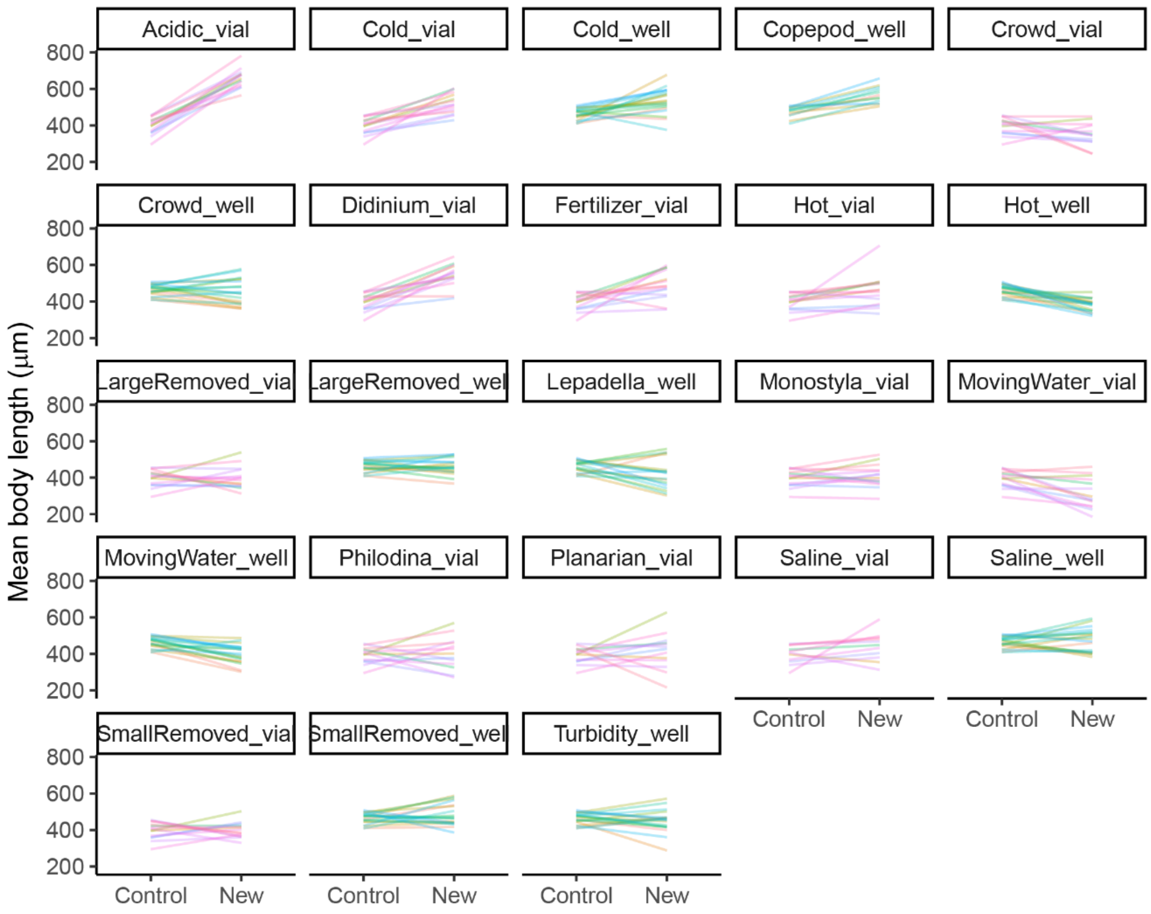


**Fig. S2. Replicate experimental populations differed in the degree of size plasticity expressed in new environments.** Clonal lines varied in plasticity of body length (LMM; $p < 0.0001$) expressed across new environments (panels). Each line represents the traits produced by a clonal line in the control environment versus a new environment. Lines are colored by clonal line and transparent to better show overlapping patterns. For the first biological replicate, on average control N = 5 rotifers x 5 replicates, new environment N = 5 rotifers x 10 replicate new environments. For the second biological replicate, on average control N = 5 rotifers x 3 replicates, new environment N = 5 rotifers x 13 replicate new environments x 3 technical replicates.

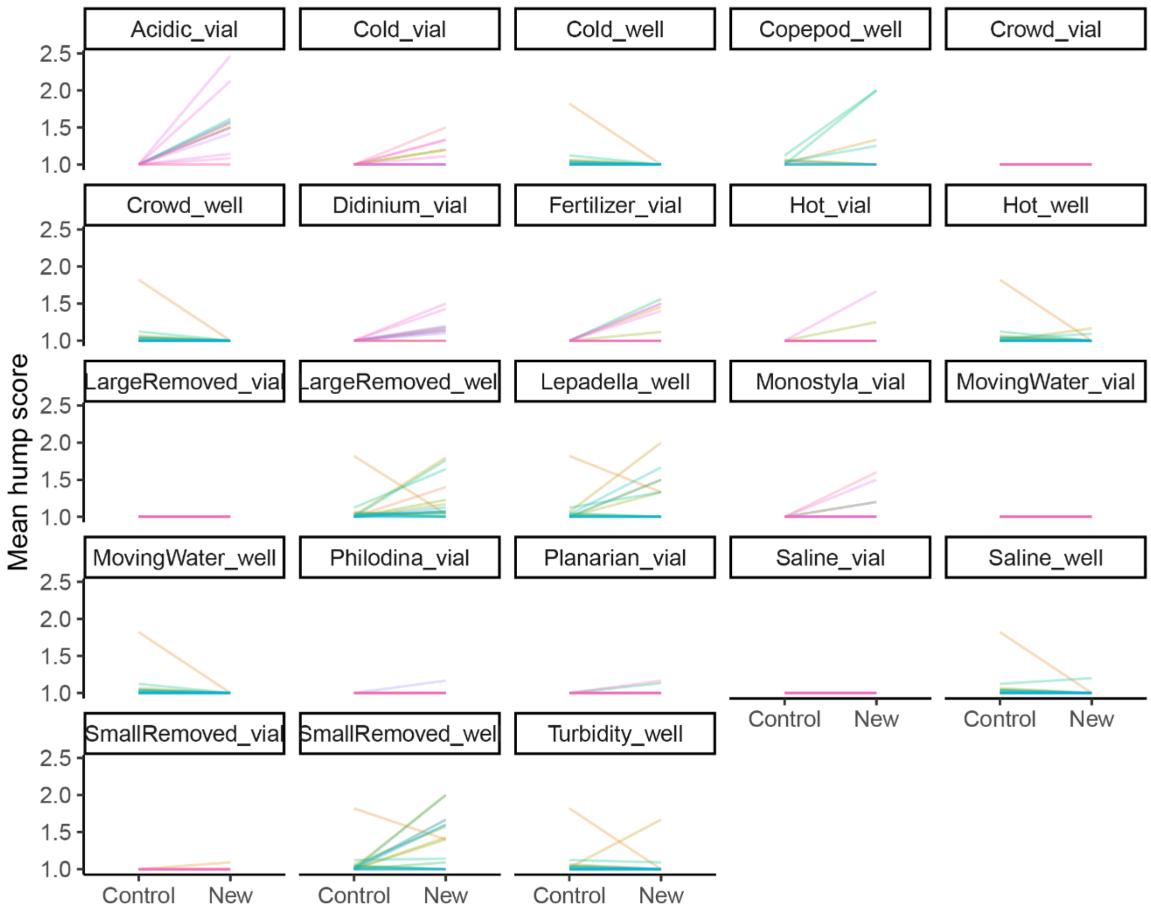


**Fig. S3. Replicate experimental populations differed in the degree of shape plasticity expressed in new environments.** Clonal lines varied in plasticity of body shape (LMM; $p < 0.0001$) expressed across new environments (panels). Each line represents the traits produced by a clonal line in the control environment versus a new environment. Lines are colored by clonal line and transparent to better show overlapping patterns. For the first biological replicate, on average control N = 5 rotifers x 5 replicates, new environment N = 5 rotifers x 10 replicate new environments. For the second biological replicate, on average control N = 5 rotifers x 3 replicates, new environment N = 5 rotifers x 13 replicate new environments x 3 technical replicates.

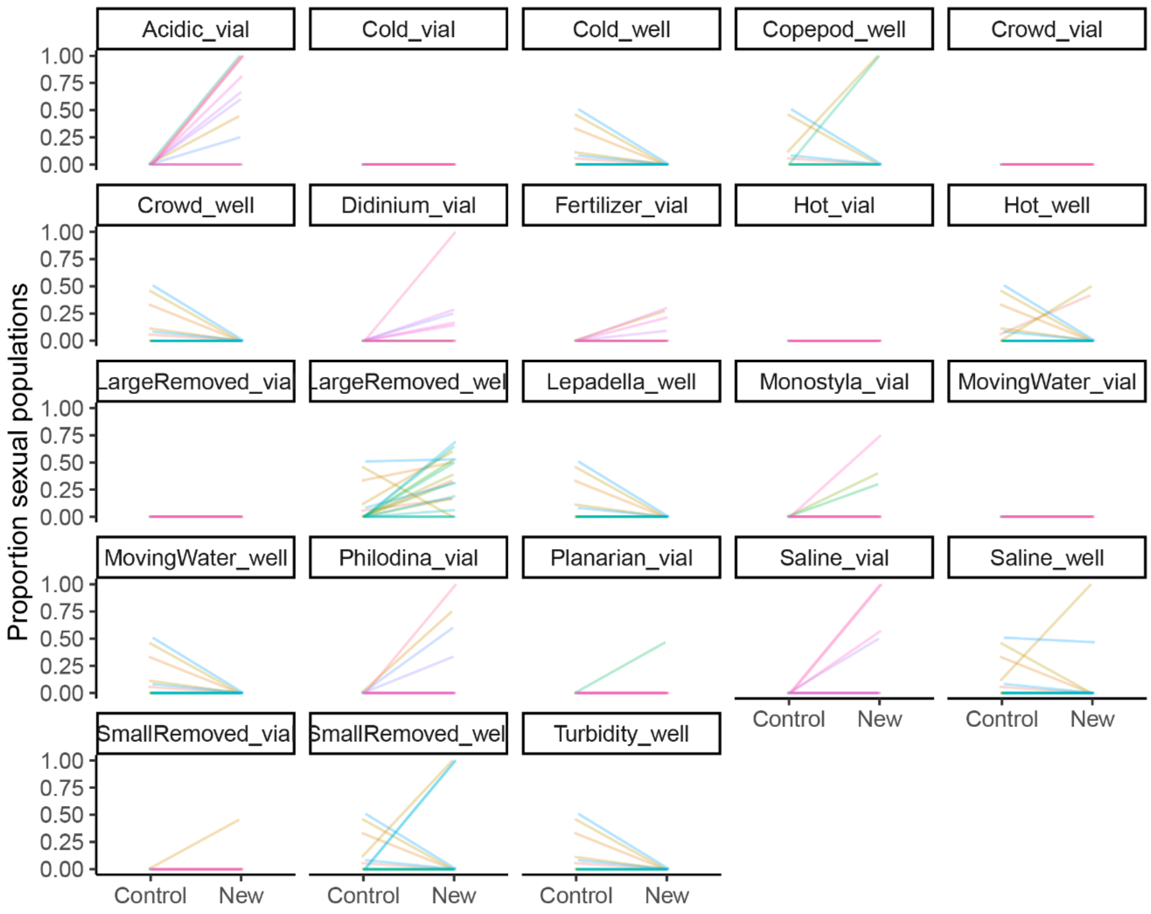


**Fig. S4. Replicate experimental populations differed in the degree of reproductive mode plasticity expressed in new environments.** Clonal lines varied in plasticity of reproductive mode (LMM; $p < 0.0001$) expressed across new environments (panels). Each line represents the traits produced by a clonal line in the control environment versus a new environment. Lines are colored by clonal line and transparent to better show overlapping patterns. For the first biological replicate, on average control N = 5 rotifers x 5 replicates, new environment N = 5 rotifers x 10 replicate new environments. For the second biological replicate, on average control N = 5 rotifers x 3 replicates, new environment N = 5 rotifers x 13 replicate new environments x 3 technical replicates.

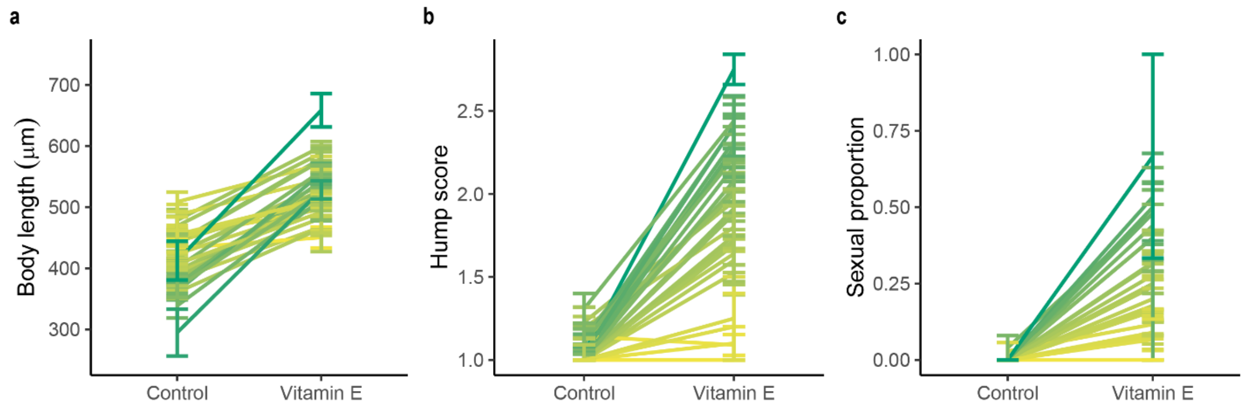


**Fig. S5. Plastic responses to vitamin E.** Reaction norms depicting the body length **(a),** body shape **(b),** and reproductive mode **(c)** a clonal line produces in the control versus vitamin E environments. Each line represents the mean value for a clonal line (on average N = 5 rotifers x 4 populations per clonal line per environment) colored on a gradient from yellow to green depicting low through high trait plasticity, respectively. Error bars ± 1 SE for replicate clones in panels a-b, ± 1 SE for replicate clonal populations in panels c.

**Table S1. Summary of differences in lifespan and fecundity between morphs.** Lifespan result is output from a linear mixed model (LMM). Test statistic is from a Kenward-Roger F test. Fecundity result is output from a negative binomial generalized linear mixed model (GLMM). Test statistic is from a likelihood ratio test. Bold values denote statistical significance at an alpha of 0.05.

| Model terms | Morph | Estimate ± SE | 95% CI | DF | Test statistic | P |
|---|---|---|---|---|---|---|
| Lifespan ~ Morph + (1\|Clonal line) | Asexual beta | -34.70 ± 9.99 | -54.05 – -14.79 | 2, 66 | 8.07 | **0.00073** |
| | Sexual beta | -39.86 ± 11.77 | -62.69 – -16.98 | | | |
| Fecundity ~ Morph + (1\|Clonal line) | Asexual beta | -0.40 ± 0.15 | -0.70 – -0.11 | 2 | 9.58 | **0.0083** |
| | Sexual beta | -0.47 ± 0.18 | -0.84 – -0.11 | | | |

**Table S2. Pairwise comparisons of morphs for lifespan and fecundity.** Pairwise comparisons of the estimated marginal means using posthoc tests with the Tukey's method for multiple comparison p-value adjustments. Test statistic is t for lifespan LMM and z for fecundity GLMM. Bold values denote statistical significance at an alpha of 0.05.

| Response variable | Morph contrast | Estimate ± SE | DF | Test statistic | P |
|---|---|---|---|---|---|
| Lifespan | Alpha – Asexual beta | 34.70 ± 10.1 | 63 | 3.44 | **0.0029** |
| | Alpha – Sexual beta | 39.86 ± 12.2 | 68 | 3.28 | **0.0047** |
| | Asexual beta – Sexual beta | 5.15 ± 12.3 | 68 | 0.42 | 0.91 |
| Fecundity | Alpha – Asexual beta | 0.40 ± 0.15 | Inf | 2.67 | **0.021** |
| | Alpha – Sexual beta | 0.47 ± 0.19 | Inf | 2.57 | **0.028** |
| | Asexual beta – Sexual beta | 0.071 ± 0.20 | Inf | 0.36 | 0.93 |

**Table S3. Variation in plasticity predicts variation in persistence.** Dormant populations are counted as extinct. Fixed effects were centered and scaled. First model is a binomial GLMM. Second model is a survival model fit as a Cox mixed model. Random effects accounting for no variance were dropped from the models. Bold values denote statistical significance at an alpha of 0.05.

| Model terms | Fixed effect | Estimate ± SE | 95% CI | DF | LRT | P |
|---|---|---|---|---|---|---|
| Persist probability ~ Size plasticity + Shape plasticity + Reproductive mode plasticity + Round + (1\|Environment) + (1\|Clone) + (1\|Rack) | Size plasticity | -0.20 ± 0.34 | -0.87 – 0.47 | 1 | 0.31 | 0.58 |
| | Shape plasticity | 0.96 ± 0.36 | 0.25 – 1.67 | 1 | 6.52 | **0.011** |
| | Reproductive mode plasticity | 0.73 ± 0.34 | 0.061 – 1.40 | 1 | 4.39 | **0.036** |
| | Round | 0.38 ± 1.82 | -3.20 – 3.95 | 1 | 0.03 | 0.85 |
| Extinction ~ Size plasticity + Shape plasticity + Reproductive mode plasticity + Round + (1\|Pond/Clonal line) + (1\|Environment) + (1\|Rack) | Size plasticity | 1.02 ± 0.13 | -0.22 – 0.27 | 1 | 0.031 | 0.86 |
| | Shape plasticity | 0.58 ± 0.17 | -0.87 – -0.22 | 1 | 10.73 | **0.0011** |
| | Reproductive mode plasticity | 0.69 ± 0.16 | -0.68 – -0.07 | 1 | 5.84 | **0.016** |
| | Round | 0.46 ± 0.88 | -2.50 – 0.95 | 1 | 0.78 | 0.38 |

**Table S4. Variation in plasticity predicts variation in persistence.** Dormant populations are counted as persisting. Fixed effects were centered and scaled. First model is a binomial GLMM. Second model is a survival model fit as a Cox mixed model. Random effects accounting for no variance were dropped from the models. Bold values denote statistical significance at an alpha of 0.05.

| Model terms | Fixed effect | Estimate ± SE | 95% CI | DF | LRT | P |
|---|---|---|---|---|---|---|
| Persist probability ~ Size plasticity + Shape plasticity + Reproductive mode plasticity + Round + (1\|Environment) + (1\|Clone) | Size plasticity | -0.045 ± 0.31 | -0.65 – 0.56 | 1 | 0.021 | 0.89 |
| | Shape plasticity | 1.18 ± 0.44 | 0.31 – 2.04 | 1 | 9.01 | **0.0027** |
| | Reproductive mode plasticity | 2.26 ± 0.52 | 1.24 – 3.28 | 1 | 34.95 | **<0.0001** |
| | Round | 1.79 ± 1.23 | -0.62 – 4.21 | 1 | 2.12 | 0.15 |
| Extinction ~ Size plasticity + Shape plasticity + Reproductive mode plasticity + Round + (1\|Pond/Clonal line) + (1\|Environment) + (1\|Rack) | Size plasticity | 0.97 ± 0.16 | -0.22 – 0.27 | 1 | 0.037 | 0.85 |
| | Shape plasticity | 0.42 ± 0.23 | -0.87 – -0.22 | 1 | 14.11 | **0.00017** |
| | Reproductive mode plasticity | 0.31 ± 0.27 | -0.68 – -0.07 | 1 | 18.66 | **<0.0001** |
| | Round | 0.30 ± 0.98 | -2.50 – 0.95 | 1 | 1.53 | 0.22 |

**Table S5. Variation in general plasticity does not predict variation in persistence.** Dormant populations are counted as extinct. Fixed effects were centered and scaled. First model is a binomial GLMM. Second model is a survival model fit as a Cox mixed model. Random effects accounting for no variance were dropped from the models. Bold values denote statistical significance at an alpha of 0.05.

| Model terms | Fixed effect | Estimate ± SE | 95% CI | DF | LRT | P |
|---|---|---|---|---|---|---|
| Persist probability ~ Size plasticity + Shape plasticity + Round + (1\|Environment) + (1\|Rack) | Size plasticity | 0.35 ± 0.54 | -0.71 – 1.4 | 1 | 0.41 | 0.52 |
| | Shape plasticity | -0.16 ± 0.31 | -0.77 – 0.45 | 1 | 0.31 | 0.58 |
| | Round | -0.19 ± 2.0 | -4.13 – 3.75 | 1 | 0.04 | 0.84 |
| Extinction ~ Size plasticity + Shape plasticity + Round + (1\|Pond/Clonal line) + (1\|Environment) + (1\|Rack) | Size plasticity | 1.07 ± 0.20 | -0.32 – 0.45 | 1 | 0.11 | 0.74 |
| | Shape plasticity | 1.06 ± 0.13 | -0.20 – -0.32 | 1 | 0.22 | 0.64 |
| | Round | 0.50 ± 0.84 | -2.35 – 0.95 | 1 | 0.69 | 0.41 |

**Table S6. Model results for analysis of persistence in new environments based on ancestral plasticity in response to vitamin E.** Fixed effects were centered and scaled. The first model is a binomial GLMM, the second is a Cox mixed model. Random effects accounting for no variance were dropped from the models. Bold values denote statistical significance at an alpha of 0.05.

| Model terms | Fixed effect | Estimate ± SE | 95% CI | DF | LRT | P |
|---|---|---|---|---|---|---|
| Persist ~ Size plasticity + Shape plasticity + Reproductive mode plasticity + Round + (1\|Environment) + (1\|Rack) | Vitamin E size plasticity | -0.45 ± 0.32 | -1.09 – 0.18 | 1 | 1.98 | 0.16 |
| | Vitamin E shape plasticity | -0.29 ± 0.34 | -0.96 – 0.38 | 1 | 0.71 | 0.40 |
| | Vitamin E proportion sexual plasticity | -0.49 ± 0.34 | -1.15 – 0.18 | 1 | 2.11 | 0.15 |
| | Round | -0.27 ± 1.96 | -4.11– 3.57 | 1 | 0.018 | 0.89 |
| Extinction ~ Size plasticity + Shape plasticity + Reproductive mode plasticity + Round + (1\|Pond/Clonal line) + (1\|Environment) + (1\|Rack) | Vitamin E size plasticity | 0.97 ± 0.12 | -0.26 – 0.20 | 1 | 0.047 | 0.83 |
| | Vitamin E shape plasticity | 0.97 ± 0.13 | -0.28 – 0.22 | 1 | 0.050 | 0.82 |
| | Vitamin E proportion sexual plasticity | 0.98 ± 0.12 | -0.25 – 0.20 | 1 | 0.045 | 0.83 |
| | Round | 0.58 ± 0.83 | -2.17 – 1.08 | 1 | 0.43 | 0.51 |

**Table S7. Model results for analysis of day 6 morphotype according to day 5 population density.** Proportion of populations with beta morphs (hump score > 1 and/or sexual) and proportion of beta populations with sexual reproduction tested with binomial GLMMs (random effects accounting for no variance were dropped from the model). Fixed effect tested with likelihood ratio tests. Bold values denote statistical significance at an alpha of 0.05.

| Model terms | Estimate ± SE | 95% CI | DF | LRT | P |
|---|---|---|---|---|---|
| Proportion beta ~ Population Size + (1\|Pond/Clonal line) + (1\|Environment) + (1\|Rack) | 0.0097 ± 0.040 | -0.068 – 0.088 | 1 | 0.059 | 0.81 |
| Proportion betas sexual ~ Population Size+ (1\|Clonal line) + (1\|Environment) | 0.021 ± 0.072 | -0.12 – 0.16 | 1 | 0.083 | 0.77 |

**Table S8. Model results for analysis of days 5-10 population growth rate according to day 6 morphotype.** Population growth rate tested with LMM against the presence of beta morphs in the population. Fixed effects tested with Kenward-Roger F test. Bold values denote statistical significance at an alpha of 0.05.

| Model terms | Estimate ± SE | 95% CI | DF | F | P |
|---|---|---|---|---|---|
| r ~ Morph + (1\|Environment) + (1\|Rack) | -0.040 ± 0.033 | -0.10 – 0.025 | 1, 195 | 1.39 | 0.24 |

**Table S9. Model results for analysis of population stability according to morphotype.** Absolute value of population growth rate at each day of measurement tested with LMM against the presence of beta morphs in the population. Fixed effects tested with Kenward-Roger F test. Bold values denote statistical significance at an alpha of 0.05.

| Model terms | Estimate ± SE | 95% CI | DF | F | P |
|---|---|---|---|---|---|
| Abs(r) ~ Morph + (1\|Environment) + (1\|Pond) | -0.015 ± 0.009 | -0.03 – 0.002 | 1, 250 | 2.84 | 0.093 |
